# Small and equipped: the rich repertoire of antibiotic resistance genes in *Candidate Phyla Radiation* genomes

**DOI:** 10.1101/2021.07.02.450847

**Authors:** Mohamad Maatouk, Ahmad Ibrahim, Jean-Marc Rolain, Vicky Merhej, Fadi Bittar

## Abstract

Microbes belonging to Candidate Phyla Radiation (CPR) have joined the tree of life as a new unique branch, thanks to the intensive application of metagenomics and advances of sequencing technologies. Despite their ultra-small size, reduced genome and metabolic pathways which mainly depend on symbiotic/exo-parasitic relationship with their bacterial host, CPR microbes are abundant and ubiquitous in almost all environments and are consequently survivors in highly competitive circumstances within microbial communities. They have been eventually identified by 16S rRNA analysis and represent more than 26% of microbial diversity. CPR microbes were able to survive in this context, although their defence mechanisms and phenotypic characteristic remain, however, poorly explored. Here, we conducted a thorough *in-silico* analysis on 4,062 CPR genomes to test whether these ultrasmall microorganisms might encode for antibiotic resistance (AR)-like enzymes. We used an adapted AR screening criteria with an exhaustive consensus database and complementary steps conferring their resistance functions. We conclude by reporting the surprising discovery of rich reservoir of divergent AR-like genes (n= 30,545 HITs, mean=7.5 HITs/genome [0-41] encoding for 89 AR enzymes, distributed across the 13 CPR phyla, and associated with 14 different chemical classes of antimicrobials. However, most HITs found (93.6%) were linked to glycopeptide, beta-lactams, macrolide-lincosamide-streptogramin, tetracycline and aminoglycoside resistance. Moreover, a distinct AR profile was discerned between the microgenomates group and Candidatus Parcubacteria, and between each of them and other CPR phyla. CPR cells seem to be active players during microbial competitive interactions and are well-equipped for the microbial combat in different habitats, supporting their natural survival/persistence and continued existence.

## Introduction

The increased use of exploring tools in the 21^st^ century, such as high-throughput sequencing and its wide application in metagenomics, has led to broadening access to genomic data of uncultured microorganisms^1^. These previously unrecognized genomes have challenged the classical view of the tree of life and have given rise to new divisions. Representatives of these divisions have been moved out of the group of undiscovered living organisms (microbial dark matter)^2^. Among these discoveries, many questions have been raised about a new group of microbes which is close to bacteria, but which is quite unique, referred to as Candidate Phyla radiation or CPR^3,4^.

CPR is a group of highly distinct and abundant ultra-small microbes, which represents more than 26% of known bacterial diversity^2^. These microbes are characterised by their reduced-size genomes^5^ and the occurrence of a high percentage of unknown-function proteins^6^. Recently, a comparative study of protein families between CPR and bacteria showed that CPR have a prevalence of proteins involved in a symbiotic lifestyle and interaction with other microbes^6,7^Therefore, they are highly auxotrophic with a lack of essential encoding genes for some pathways which are critical to the autonomous lifestyle^8^.

Paradoxically, the lack of these genes can sometimes help them to survive in their habitat. For example, despite the absence of a viral CRISPR defence system in *Patescibacteria* (the phylum that contains most CPR genomes), members of this superphylum can escape bacteriophage attacks (attachment) by the natural suppression of common phage membrane receptors^9^.

However, these as yet uncultured microbes have been detected based on metagenomic or metabarcoding analyses of ribosomal RNA sequences^3^. To date, CPR microbes have been reported in different human microbiomes (buccal cavity, gut microbiota, vagina etc.)^10,11,12,13,14^, as well as in the environment (soil, seawater, deep-sea sediments, termite guts etc.)^15,16,17,18,19,20^. Their ubiquitous presence in complex ecosystems therefore suggests their continuous competitive lifestyle against different microorganisms. This focusses attention on understanding the defensive mechanisms employed by CPR microbes in habitats shared with other microbes.

Moreover, according to metagenomic analyses of ancient DNA, CPR microbes have been reported in ancient samples of Neanderthal calcified dental plaque (calculus) dated thousands of years ago^21^. Like CPR, antibiotic resistance (AR) is an ancient phenomenon highly reported in the microbial world^22,23^. Various studies have shown the natural existence of AR genes in micro-organisms even before the discovery and introduction of antibiotics by humans in the mid-twentieth century^24^. These AR genes have also been detected from ancient samples dating back millions of years in diverse environments^24^. The mechanisms of AR are due to the absence of antibiotic targets, their modification following a mutation on pre-existing genes, or to the presence of protein coding genes^25^. Some genes can inactivate the antibiotic by enzymatic activity, while other genes confer AR by target protection or alteration^25^.

Given that CPR members (i) are widely spread in different ecological niches and microbiomes, (ii) have never been isolated and grown in pure culture, and (iii) have a high number of unknown biosynthetic activities within their genomes, few, if any studies have looked into the defence mechanisms and competing behaviour of CPR cells. In fact, survival strategies, which are pointedly AR gene components expressed by CPR members against other microbes in different hostile/competitive environments, have not yet been explored. For this propose, we describe the first repertoire of AR genes in CPR genomes by *in silico* analysis, after developing suitable AR screening criteria. We found that CPR members are also players in this microbial “infinity war”.

## Materials and Methods

### Genomic data

For this study, all nucleotide sequences of CPR genomes available on 12 September 2020 on the NCBI website (National Center for Biotechnology Information) (https://www.ncbi.nlm.nih.gov) were selected and downloaded from the NCBI-GenBank database. Genomes were chosen based on the taxonomy provided by the NCBI. The 4,062 CPR genomes are distributed across 2,222 Candidatus Parcubacteria, 933 Candidatus Microgenomates, 284 Candidatus Saccharibacteria, 155 unclassified Patescibacteria group, 136 Candidate division WWE3 (Katanobacteria), 126 Candidatus Peregrinibacteria, 55 Candidatus Berkelbacteria, 53 Candidatus Dojkabacteria, 39 Candidatus Doudnabacteria, 33 Candidatus Gracilibacteria, 13 Candidatus Absconditabacteria, 11 Candidate division Kazan-3B-28 and two Candidatus Wirthbacteria. Only 35 of all the genomes analysed were complete genomes, while the remaining were whole genome sequences (WGS).

Genome annotation was generated using the Rapid Annotation using Subsystem Technology tool kit (RASTtk)^26^ as implemented in the PATRIC v3.6.8 annotation web service.

### Detection of Antibiotic Resistant Genes in CPR genomes

For antimicrobial resistance profiling, we carried out an in-house Blast search against the protein databases from ARG-ANNOT (Antibiotic Resistance Gene-ANNOTation)^27^, BLDB (Beta-Lactamase DataBase)^28^ and NDARO (National Database of Antibiotic Resistant Organisms)^29^ containing 2,038, 4,260 and 5,735 sequences, respectively. In order to get a comprehensive view of the CPR resistome we used relaxed parameters including a minimum percent of identity and coverage length equal to 20% and 40%, respectively, and a maximum E-value of 0.0001^6^. All results were checked manually to remove duplications.

Predicted ARs in each CPR genome were individually compared to proteins in each AR database by reciprocal BLASTP^30^. The number of reciprocal best hits was counted using an expectation value (E) of 0.0001 as the stringency threshold for determining a valid best hit. Only the CPR protein sequence resulting from the reciprocal BLASTp and matched with the same AR gene resulting from the first BLASTp was conserved for the next step as the preliminary results of AR genes.

In order to eliminate false positive HITs, a BLASTp search of the preliminary AR genes as a query data set was performed against the conserved domains database (CDD) (https://www.ncbi.nlm.nih.gov/Structure/cdd/wrpsb.cgi). The AR predicted genes with a protein domain necessary for the AR mechanism were subsequently selected. A literature review was conducted for each family of antibiotics detected in the CPR genomes to determine the mechanism of AR. We were only interested in the genes in which the AR mechanism depends on enzymatic activity and didn’t consider the mechanisms that require a further search for site mutations (Figure 1).

**Figure 1:**
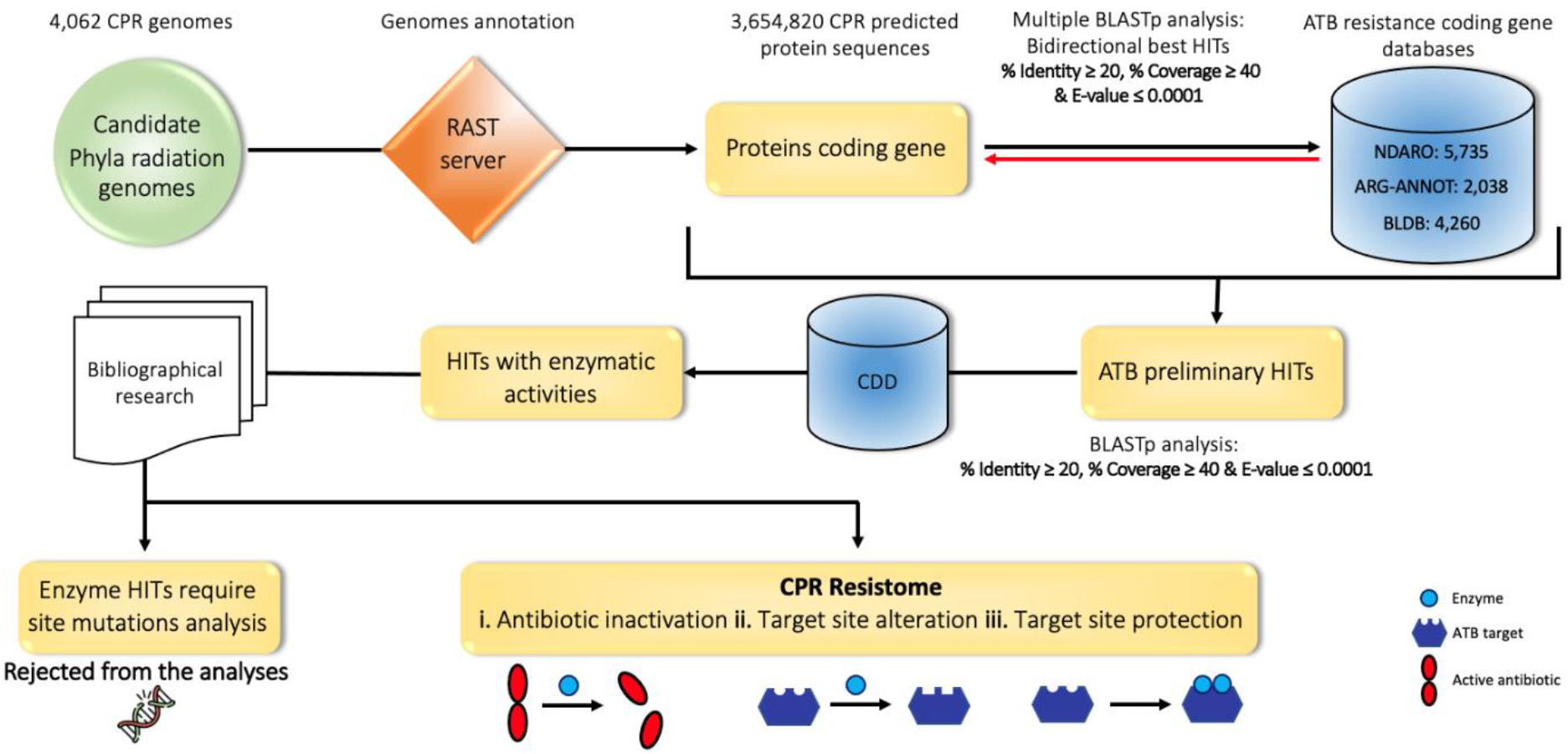
Study design. The first step consists of annotating the CPR genomes available on the NCBI website, using the RAST server. The CPR protein sequences are considered as queries for BLASTp against consensus databases of bacterial antibiotic resistance (AR) genes. The analysis was performed with a minimum identity and coverage percentage of 20% and 40%, respectively, and a maximum E-value of 0.0001. The AR preliminary HITs resulting from the simple BLASTp are queried against the multiple databases of AR genes as performing a reciprocal BLASTp. Further analyses were undertaken to detect the protein functional domain for HITs with enzymatic activity using the conserved domain database (CDD). Finally, bibliographical research was conducted to select enzymes conferring resistance with specific mechanisms of actions as CPR resistome.

AR-like genes detected in CPR tested genomes are represented using Cytoscape v.3.8.2 to highlight the link between different antibiotic families and distinct CPR phyla. These genes are also represented in a multi-informative heat map performed by Displayr online tool (www.displayr.com), to show the distribution of different AR-like genes on CPR phyla and their mechanisms of AR.

## Results

### CPR microbes encode for vastly divergent AR-like genes according to reference bacterial protein databases

In this study, we adapted a suitable strategy for the specific detection of AR like genes in the 4,062 CPR genomes tested. The simple BLASTp of the 3,654,820 CPR protein sequences predicted from the coding DNA sequences (CDS) detected using the RAST server, against a total of 12,033 AR protein sequences resulted in 320,121 HITs. After performing the reciprocal BLASTp search, our analyses led to 175,238 preliminary AR HITs with the conservation of the protein functional domains necessary for the resistance mechanisms, as mentioned above (see also materials and methods). We then focused only on enzyme encoding genes that confer resistance to a given antibiotic family. However, after eliminating all HITs corresponding to mutations (134,693 HITs), we retained a total of 30,545 HITs, corresponding to a total of 89 AR-like genes for further analyses (Figure 2 and Table S1). These genes constituted the target data set in our analysis and were considered as the CPR resistome. This is used for deciphering the high potential of proto-resistance genes as a deep reservoir of AR in these micro-organisms.

**Figure 2:**
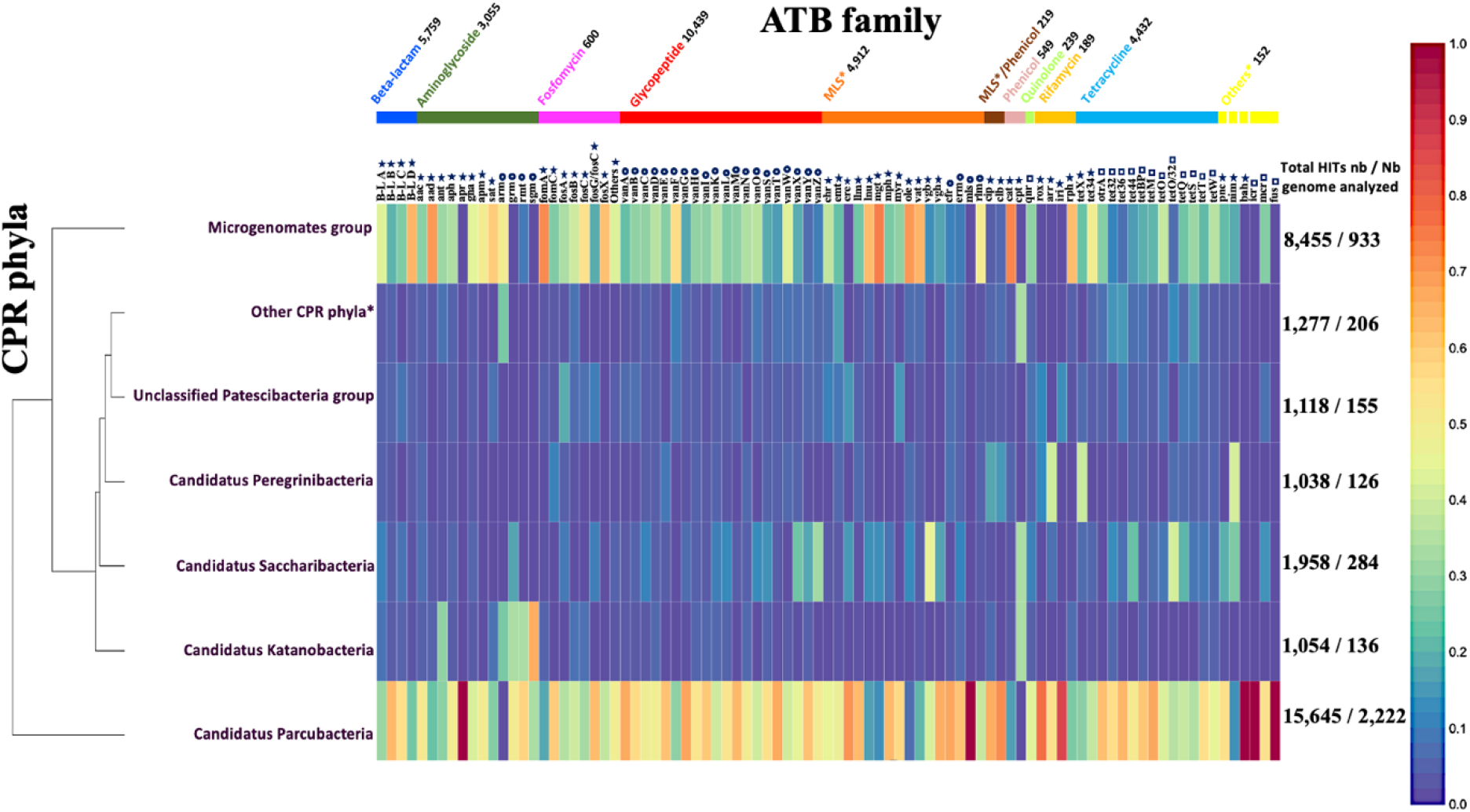
Multi-informative heat map of antibiotic resistance (AR) like genes in CPR genomes. Detection of 30,545 AR-like genes in 4,062 CPR genomes using an adapted AR screening strategy. The abundance of each AR-like gene on each CPR phylum is relative to the total number of AR-like genes found in all CPR phyla (Number of AR-like genes found in CPR phylum divided by the total HITs number of this AR family). MLS* indicates the merging of the three antibiotic families: macrolide, lincosamide and streptogramin. Others* indicates the merging of five antibiotic families with fewer AR-like genes: pyrazinamide, nitroimidazole, bacitracin, colistin and fusidic acid. **✶** indicates AR-like genes that confer resistance by antibiotic inactivating enzymes, o indicates AR-like genes that confer resistance by antibiotic target alteration and **□** indicates AR-like genes that confer resistance by antibiotic target protection. The other CPR phyla* indicate the merging of all Candidatus CPR phyla with fewer than 100 genomes: Candidatus Berkelbacteria, Candidatus Doudnabacteria, Candidatus Wirthbacteria, Candidate division Kazan, Candidatus Dojkabacteria, Candidatus Absconditabacteria and Candidatus Gracilibacteria.

Most AR HITs found in CPR had a similarity percentage ranging from 30% to 40% against bacterial AR genes (Figure 3), highlighting the divergence of their sequences from those of bacteria. These findings support the prediction of resistance enzymes encoding genes in CPR microbes, but also suggest that these enzymes may differ slightly from well-characterised bacterial ones.

**Figure 3:**
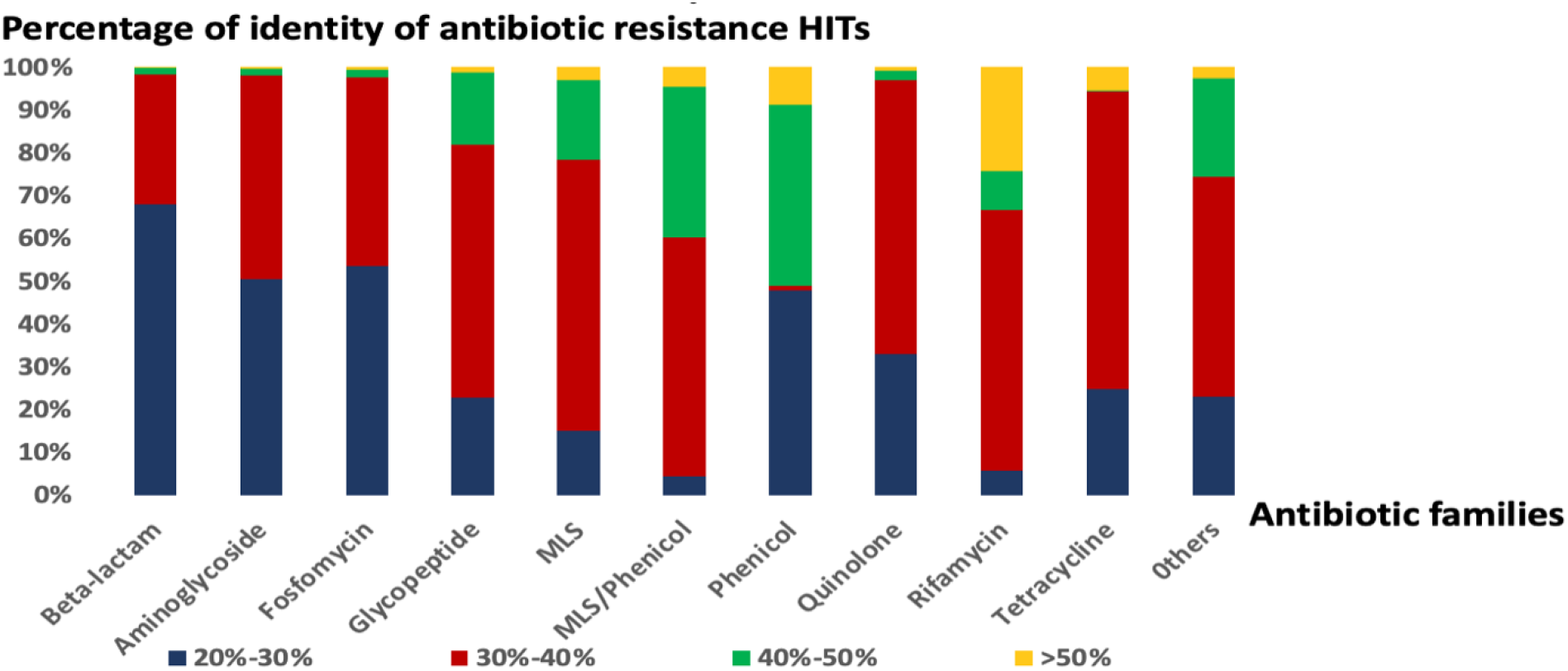
Histogram representing the distribution of antibiotic resistant (AR) HITs by percentage of similarity against bacterial AR genes in each antibiotic family detected in this study. The others indicate the merging of four antibiotic families with fewer AR-like genes: pyrazinamide, nitroimidazole, bacitracin, colistin and fusidic acid.

The diversity of the CPR resistome involves 14 different antibiotic families: 34.18% glycopeptide, 18.85% beta-lactam, 10% aminoglycoside, 14.51% tetracycline, 16.08% MLS for macrolide-lincosamide-streptogramin, 1.8% phenicol, 1.96% fosfomycin, 0.62% rifamycin, 0.78% quinolone and 0.5% of other antibiotic families (bacitracin, fusidic acid, pyrazinamide, nitroimidazole and lipopeptides) (Figures 2 and 4 and Table S1). A high percentage of the AR HITs identified in our study confer AR by altering its target, with methyltransferase activity of 16S ribosomal RNA (47.34% of total HITs: 14,459 HITs), whereas others act directly on a given antibiotic by inactivating it (37.73% of total HITs: 11,525 HITs) or by protecting its target (14.93% of total HITs: 4,561 HITs) (Figures 2 and 4 and Table S1).

**Figure 4:**
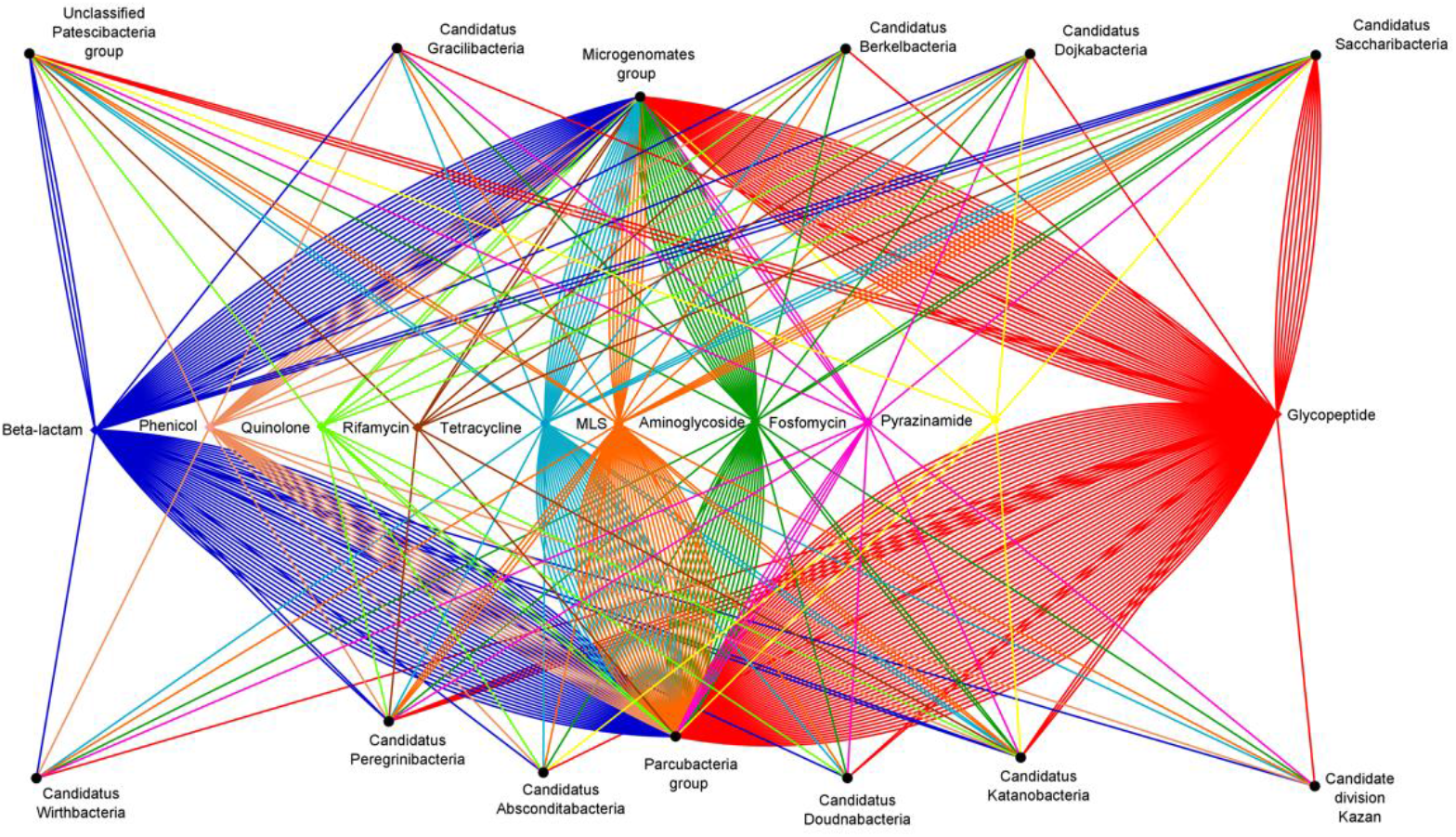
Network analysis of antibiotic resistance-like gene distribution, highlighting the link between different antibiotic families and distinct CPR phyla. Beta-lactam in blue, phenicol in rose, quinolone in light green, rifamycin in brown, tetracycline in light blue, MLS in orange, aminoglycoside in dark green, others including pyrazinamide, nitroimidazole, bacitracin, colistin and fusidic acid in yellow, fosfomycin in pink and glycopeptide in red. An asterisk (*) indicates the merging of the three antibiotic families: macrolide, lincosamide and streptogramin into one MLS family.

Equally, we found AR HITs in almost all CPR genomes which we tested across different phyla; 4,052 genomes were positive through our analysis out of 4,062 genomes tested (99.75%). The prevalence of the AR content is fairly diversified between the CPR phyla, as the number of their available genomes is not homogeneous (Figures 2 and 4 and Table S1). Furthermore, each CPR phylum holds at least resistances to six different classes of antimicrobials, and they have nearly the same distribution of AR HITs (Figure 5). The resistance to common antibiotic families found in different CPR phyla represent five of the total families identified, namely glycopeptide, beta-lactam, MLS, tetracycline, and aminoglycoside, highlighting the importance of the function of these AR HITs in CPR genomes.

**Figure 5:**
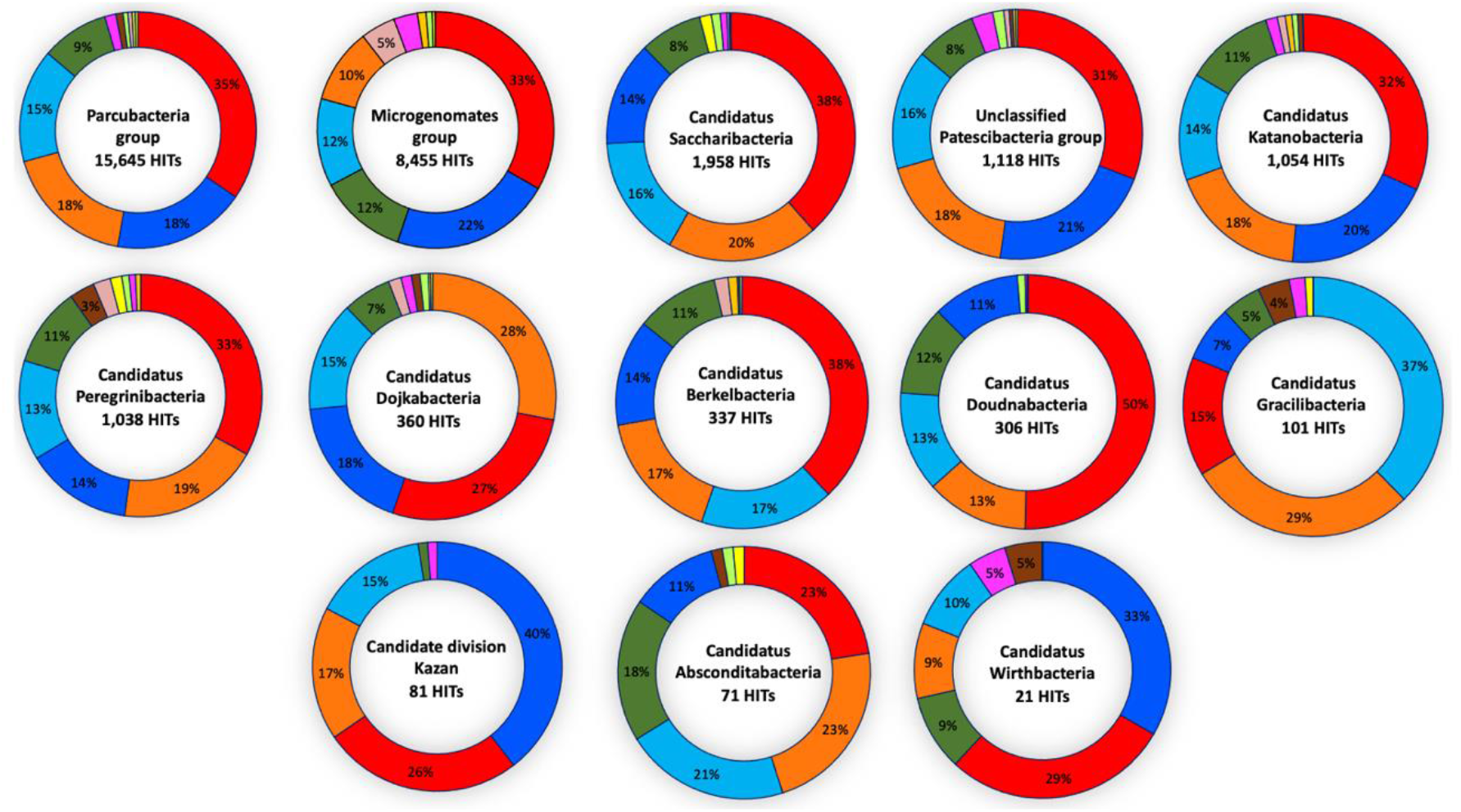
The distribution of the percentage of antibiotic resistance-like genes by antibiotic families on each CPR phylum. Glycopeptide (red), beta-lactam (blue), aminoglycoside (dark green), tetracycline (light blue), MLS (orange), phenicol (rose), fosfomycin (pink), rifamycin in gold, MLS/phenicol (brown), quinolone (light green), pyrazinamide (yellow), nitroimidazole (yellow), bacitracin (yellow), colistin (yellow) and fusidic acid (yellow).

### The prevalence of detected enzymes according to each chemical antimicrobial class

In this part, we took all chemical classes of antimicrobials for which we could detect HITs in all CPR phyla. Starting with glycopeptide, the resistance hits were found to be the most abundant AR-like genes. This resistance involving vancomycin susceptibility is caused by a modification of the antibiotic target D-Alanine:D-Alanine into D-Alanine:D-Lactate or D-Alanine:D-Serine. Since vancomycin resistance is mediated by a cluster of genes including essential, regulatory, and accessory genes, we searched only for CPR genomes with the three essential genes in the cluster. These can be classified into nine types based on their genetic sequences and structures: vanA, vanB, vanC, vanD, vanE, vanG, vanL, vanM and vanN.

Forty-eight of the CPR genomes have a potential for vancomycin resistance as they carry the three essential genes for the functioning of a given cluster. We looked for the gene that gives the cluster name, plus vanH and vanX for D-Ala:D-Lac clusters and vanX and vanT for D-Ala:D-Ser clusters. We found a total of 18 D-Ala:D-Lac vancomycin clusters, including nine vanA clusters, three vanB clusters, five vanD clusters and one vanM cluster. For D-Ala:D-Ser ligase gene clusters, we found 25 vanC clusters, one vanE cluster, three vanG clusters, three vanL clusters and two vanN clusters (a total of 34 D-Ala:D-Serine vancomycin clusters) (Figure S1). Of these 48 genomes, four had two different types of vancomycin clusters: one genome presented the essential genes of the vanB and vanD clusters, one genome with vanA and vanC clusters and two genomes with vanL and vanN clusters (Figure S1). More analysis is needed to search for the presence of other components (regulatory and accessory genes) and the synteny of these genes as they participate together in the correct functioning of the vancomycin cluster.

Given that CPR members have very small genomes in comparison with other microorganisms, it is profitable for these microbes to have multifunctional genes, such as beta-lactamases. The 5,759 beta-lactam resistant HITs belong to four different classes (A, B, C and D) (Figure 4). Class B metallo-beta-lactamases are the most frequent, representing 58.3% of those detected (3,359 HITs over 5,759 HITs) (Table S1). This class has been classified into three different subclasses of metallo-β-lactamases depending on the annotation of the CDD results: 17 HITs belong to subclass B1, 385 HITs to subclass B2 and 2,957 HITs to subclass B3. Moreover, 2,400 of the serine-beta-lactamases are distributed over 27.9% of class A, 0.5% of class C and 13.3% of class D (Table S1).

For macrolide-lincosamide-streptogramin (MLS), the most common genes (erm (n=3,077 HITs) and cfr (n=648 HITs)) (Table S1), detected in the CPR genomes, are involved in MLS resistance by altering its target with esterase activity and methylation of the 23S rRNA subunit, respectively, followed by streptogramin acetyltransferase (vat; n=338 HITs) (Table S1) with MLS inactivating enzyme activity. In addition, our study showed aminoglycoside resistance HITs in all CPR phyla with different transferase activities: adenylyltransferase, phosphotransferase, and acetyltransferase. The majority code for acetyltransferase activity, of which the most abundant genes are aac (aminoglycoside acetyltransferase, n= 1,831) and gna (gentamicin acetyltransferase, n=749) (Table S1). Finally, almost all tetracycline resistant HITs confer resistance through ribosomal protection and code for tetT (n= 2,243), tetBP (n= 778) and tetW (n= 564) (tetracycline resistance ribosomal protection protein) (Table S1).

### Antibiotic resistance profile according to CPR phyla

Based on our AR screening strategy, only 10 genomes were found to be negative from a total of 4,062 CPR genomes analysed. The others were found to be positive, with a notable average of 7.5 AR-like genes per genome. The general distribution of HITs classed according to antibiotic family was almost maintained in the various CPR phyla, with some exceptions (Figure 5), despite the high difference in the number of AR HITs found between the Parcubacteria phylum regrouping most CPR genomes and Candidatus Wirthbacteria (15,645 AR HITs in 2,222 tested genomes compared to 21 AR HITs in two genomes, respectively) (Figure 4 and Table S1). Interestingly, CPR phyla were clustered into three major groups according to their AR content and the abundance of the detected genes (Figure 4 and Figure S2). The first group includes Parcubacteria genomes, the second includes Microgenomates genomes, and the last group includes the remaining CPR phyla. Three different AR profiles were therefore identified for CPR phyla. In the microgenomates group, we observed a significant number of genes with adenylyltransferase (aad) and acetyltransferase (gna) activity against aminoglycosides, phosphorylation of fosfomycin (fomA) and a remarkable number of class A and D beta lactamases (Table S1). Moreover, this group of microgenomates possess the greatest number of cat (chloramphenicol acetyltransferase) enzyme encoding genes, streptogramin acetyltransferase (vat) and rifampin phosphotransferase (rph) detected among all CPR genomes. In contrast, taking Saccharibacteria as an example of the group of other CPR phyla, members of this phylum have a high number of streptogramin lyases (vgb) and erythromycin esterases (erm) compared to other CPR groups (Figure 2 and Table S1).

It should be noted that the more available genomes we analysed, the more likely we were to detect additional antibiotic families. This is the case for the Parcubacteria group, where bah (the amidohydrolase enzyme that inactivates bacitracin), icr (intrinsic colistin resistance enzyme) and fus (fusidic acid resistance enzyme) were found only in this CPR group (Figure 2 and Table S1).

To sum up, these results are suggestive of the influence of the CPR environment on its phenotypic characteristics and suggest a link between CPR members, other microbes, and their environment. Together, the high presence of AR-like genes in all CPR genomes indicates that they are likely to be functionally linked to other metabolic pathways and, subsequently, to participate in the survival of these microorganisms.

## Discussion

There are significant knowledge gaps in our understanding of the physiological and biological processes of CPR, as well as of their interactions with host bacteria and their potential associations with human pathologies. Thus, it is essential to expand our research on these living microorganisms, which represent a new branch in the tree of life^31^. This study aimed to report the existence of AR in these ultra-microbes and to determine the AR profile of each CPR phylum. These analyses may contribute towards a better elucidation of CPR phenotypic characteristics and its defence mechanisms.

In our study, we conducted thorough *in-silico* screening for AR in all CPR genomes. Our analysis was based on an adapted strategy for this new branch of the tree of life, using multiple computational methods. We revealed a rich repertoire of AR genes encoded by almost all tested CPR genomes. We allocated the AR-like genes into distinct approaches in order to visualise the prevalence of AR genes in different CPR phyla and, potentially, to find a correlation between resistance genes to a particular antibiotic family and the phylum of interest.

Since resistance has never been searched for in CPR before and given that CPR microbes have not yet been grown in pure culture, their resistance can only be explored by *in-silico* analysis for the moment. AR screening of CPR genomes by analysing nucleotide sequences against a database of bacterial resistance genes (the classical method of AR profiling in the bacterial domain)^27^ resulted in a negligible number of HITs when compared with our optimised strategy (data not shown).

It was critical to establish an adapted strategy for AR screening in CPR genomes, as they have original nucleotide and protein sequences^1^. Based on the evolutionary variation of sequences, the protein sequences involved in the biological function proceed at a slow rate, unlike those of nucleotides^32^. We therefore used protein sequences in our strategy. The genomes were annotated by using the RAST server, as it had the lowest percentage of unannotated proteins, despite giving a high percentage of hypothetical proteins^33^, which is standard in CPR, as high numbers of their metabolic pathways and biosynthetic capacities have not yet been determined^7^.

Attempting to study a new branch of the tree of life when there is a huge lack of data is challenging. Hence, it relies on previously known information. For this reason, less stringent parameters were used to achieve a more comprehensive exploration of the AR contents^6^. Moreover, we use multiple AR gene databases to detect maximum HITs, since there is currently no specific AR database for CPR members. However, a reciprocal BLASTp was performed to reduce the number of false positive results. The functional protein domains were then searched for against the detected HITs, as its essential to retain all necessary patterns related directly to the biological function of these sequences. For more accurate results, our analysis only took into consideration HITs with enzymatic activities. These enzymes confer resistance by acting directly on the inactivation of the corresponding antibiotic or by its target protection or alteration. AR HITs with mutations were discarded from further analyses since CPR sequences are not comparable with or similar to those of bacteria. Our multistep study design guarantees an optimal balance between the intended function (specificity) and permissive stringency (sensitivity).

Nevertheless, this strategy may also miss some resistance genes and thus lead to false negative results. It could be expected that CPR members have antimicrobial resistance sequences that are significantly different from those of bacteria, with new patterns and undescribed resistance mechanisms, particularly because CPR microbes have divergent sequences from those of bacteria due to rapid evolutionary phenomena^1^. In addition to the resistance profiling found in this study, the possible presence of efflux pumps in CPR cells, as in all living microorganisms, which participate in the detoxification process by expelling various harmful and xenobiotics compounds should not be overlooked. In particular, these include the multi-drug efflux mechanisms which are normally encoded by the chromosome^34^.

The surprising and somewhat paradoxical presence of resistant genes in microorganisms with reduced genomes, such as those of CPR, raises the questions of their origin and their indispensable function. We believe that they are ancestral, due to their divergent sequences from other microbial domains of life. The transmission of these genes therefore occurs mainly through vertical gene transfers. These HITs may have other functions that are involved in different metabolic pathways rather than resistance to antibiotics.

For the resistance to the glycopeptide family, we expected our results to show a significant number of vancomycin resistance-like genes, as the function of this resistance depends on the presence of an operon of seven genes^25^. Given the significant diversity of these genes that has previously been described^35^, we detected 20 different types of vancomycin, namely vanA, vanB, vanC, vanD, vanE, vanF, vanG, vanH, vanI, vanK, vanL, vanM, vanN, vanO, vanS, vanT, vanW, vanX, vanY, and vanZ in the 4,062 CPR genomes. Additional analyses enabled us to identify 48 CPR genomes with a potential for a vancomycin resistance. These genomes feature the three essential components of a functional vancomycin resistance cluster. Further analysis of the remaining elements is required to have a complete cluster with accessory and regulatory genes. In addition to the presence of these elements, it is necessary to verify their adequate arrangement to ensure the correct functioning *in-vivo*.

However, as described previously, the membrane of CPR cells is very similar to that of Gram positive^6^ bacteria, which develop resistance to vancomycin by modifying the D-Alanine:D-Alanine peptidoglycan precursor^25^. CPR microbes may have a natural presence of regulatory genes in their genomes, including efflux pumps which were subsequently excluded from our assays (for example, vanR is present in all CPR genomes (100%) (Data not shown)). These genomes may naturally produce the D-Ala:D-Lac or D-Ala:D-Ser peptidoglycan precursors rather than the natural precursor D-Ala:D-Ala in bacteria. This supports the intelligent way in which these microorganisms survive with a limited number of genes; that is an incomplete but functional cluster (i.e., no need for accessory genes, as their names indicates). This supports the idea that the CPR genome is simple but efficient. Further analysis should be carried out to verify the AR conferred by the absence or modification of its targets, in addition to that conferred by the presence of active enzymes carried out as part of this study.

Our results also show that there is almost one beta-lactam resistant gene per CPR genome; 77% of the tested genomes have at least one gene which codes for beta-lactamases (classes A, B, C and D). These genes may play a role in the degradation of substances used in metabolic pathways, including beta-lactams. Several studies have shown that beta-lactamases are multifunctional genes which play several roles including, but not limited to, endonuclease, exonuclease, ribonuclease and hydrolase^36^. Furthermore, beta-lactamases have been detected in other life domains including bacteria^37^, eukaryotes^38^, and archaea^39^, and this may therefore also be the case for CPR. It is very likely that the presence of multifunctional genes is necessary and indispensable in CPR members, due to their small genomes and the very reduced number of genes per genome compared to other microorganisms.

Interestingly, aminoglycoside resistance has been mentioned and used for the co-culture of TM7x with its host species bacteria, *Actinomyces odontolyticus* strain XH001. The authors enriched TM7x through streptomycin selection, as its host is also highly resistant to streptomycin. It is likely that CPR members are resistant to aminoglycosides and other antibiotics targeting RNA. Besides having an uncommon ribosome composition/sequence, some CPR have introns in their 16S, 23S rRNA and tRNA^3^. Given their tiny genomes, this is a prominent feature for them to encode multifunctional genes, depending on the intron splitting.

The significant prevalence of AR genes in this new branch of the tree of life sheds light on the problem of choosing the appropriate treatment in the clinical field. It is important to investigate whether the failure of antibiotic treatment in different cases is due to the presence of hidden resistance genes or the presence of resistance genes that have not been searched for (our study has already confirmed that CPR genomes can act as resistance vectors). The failure to provide adequate treatment is related to overlooking AR screening in CPR, which may be responsible for the transfer of the AR profile to the host bacteria without gene transfer.

Finally, the AR-like genes detected in CPR genomes in our *in-silico* screening are expected to be confirmed in upcoming *in-vitro* experiments. A specific database for AR gene screening in CPR genomes needs to be created to collect these new results for further studies.

## Conclusion

This work contributes towards a new way of deciphering this new branch of the tree of life. We explicitly explored the CPR resistome by establishing an adapted AR screening strategy for these fastidious micro-organisms. We found a gigantic reservoir of AR, representing the first report of resistance genes in CPR genomes. These highly abundant microbes could be an interesting paradigm which constitutes an endless natural source of emerging resistances. Our findings represent a substantial opportunity for future scientific discoveries. If, as expected, the AR-like genes detected in CPR are involved in different metabolic pathways, further studies may lead to of the successful growth of CPR cells in pure culture.

## Funding information

This work was supported by the French Government under the “Investissements d’avenir” (Investments for the Future) programme managed by the Agence Nationale de la Recherche (ANR, fr: National Agency for Research), (reference: Méditerranée Infection 10-IAHU-03).

This work was supported by Région Provence Alpes Côte d’Azur and European funding (FEDER (Fonds européen de développement régional) PRIMMI (Plateformes de Recherche et d’Innovation Mutualisées Méditerranée Infection)).

## Conflicts of interest

The authors declare that they have no competing interests.

---

**Figure S1:**
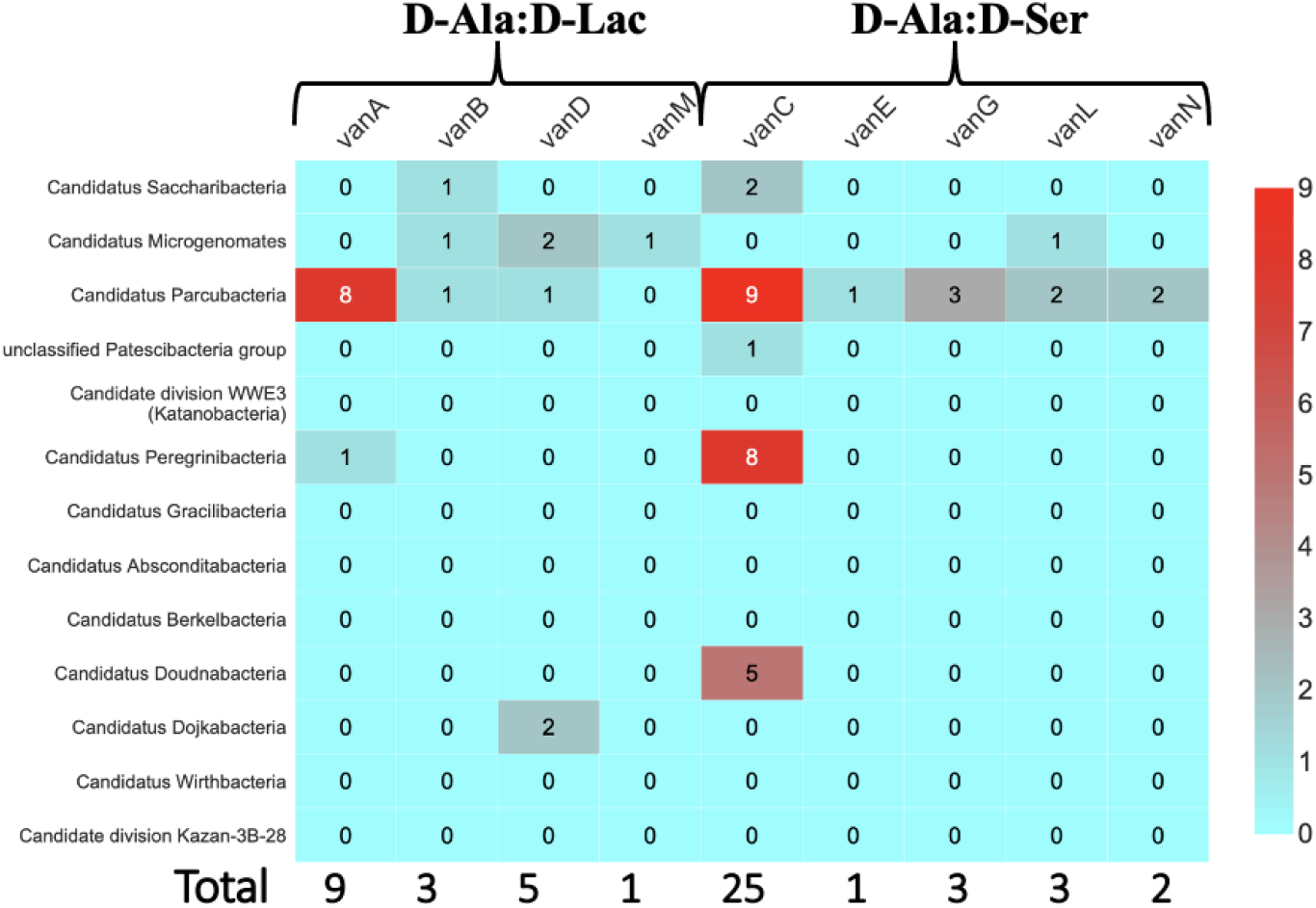
Heat map of different vancomycin clusters D-Alanine:D-Lactate and D-Alanine:D-Serine with the presence of the three essential genes in the CPR phyla tested. 48 CPR genomes have a total of 52 potential of vancomycin clusters including four genomes each with two types of clusters.

**Figure S2:**
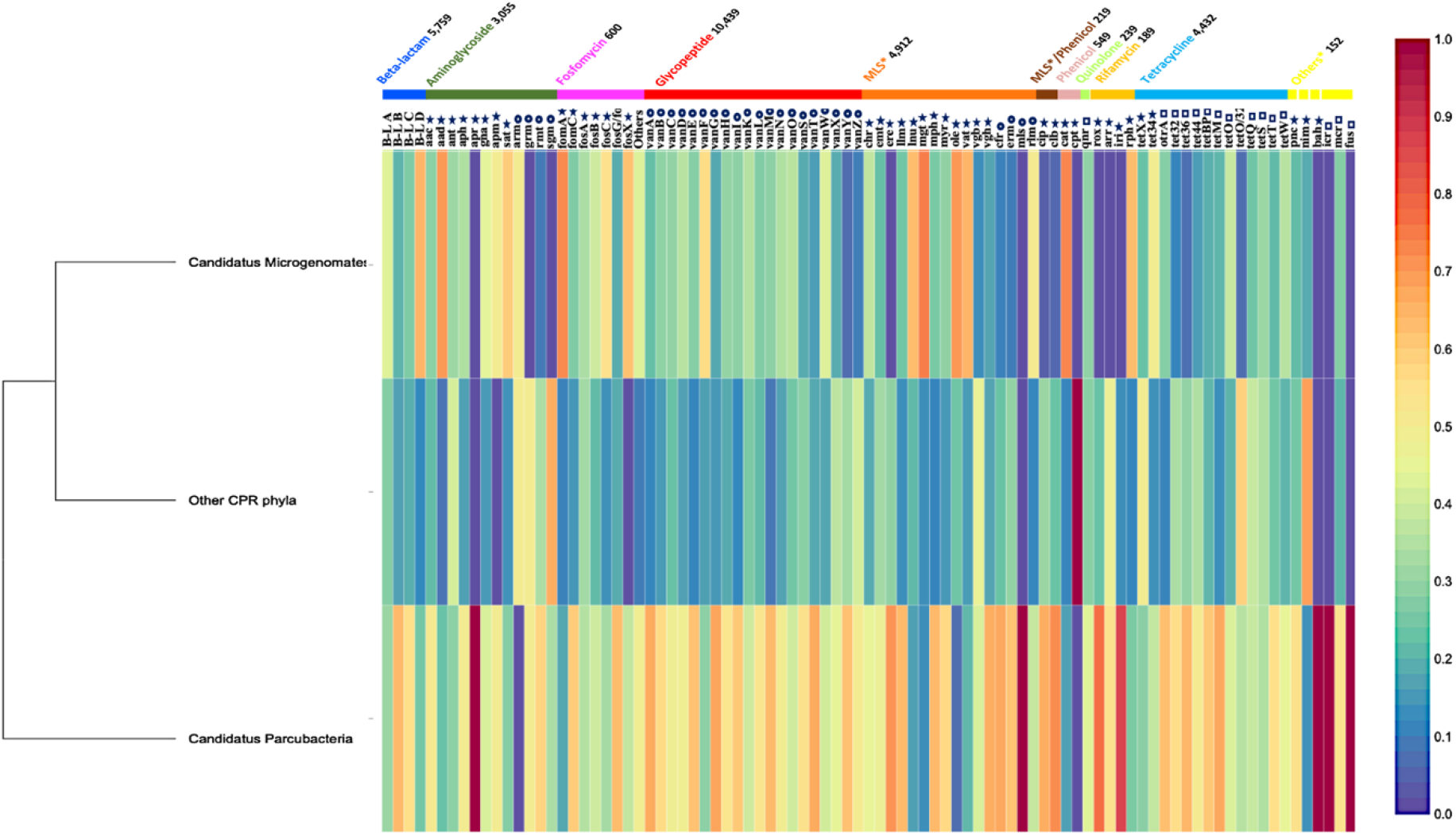
Multi-informative heat map (Figure 4) with the grouping of Saccharibacteria, unclassified Patescibacteria group, Katanobacteria, Peregrinibacteria, Berkelbacteria, Dojkabacteria, Doudnabacteria, Gracilibacteria, Absconditabacteria, Kazan-3B-28 and Wirthbacteria together as the other CPR group.

## Notes

### Competing Interest Statement

The authors have declared no competing interest.

