## Supplementary material for "Small and equipped: the rich repertoire of antibiotic resistance genes in *Candidate Phyla Radiation* genomes": Table S1

Table S1 : number of AR HITs found according to CPR Phyla

| Gene | Abbreviation | Candidatus Saccharibacteria | Candidatus Microgenomates | Candidatus Parcubacteria | unclassified Patescibacteria group | Candidate division WWE3 (Katanobacteria) | Candidatus Peregrinibacteria | Candidatus Gracilibacteria | Candidatus Absconditabacteria | Candidatus Berkelbacteria | Candidatus Doudnabacteria | Candidatus Dojkabacteria | Candidatus Wirthbacteria | Candidate division Kazan-3B-2 | Total |
|---|---|---|---|---|---|---|---|---|---|---|---|---|---|---|---|
| Beta-lactam classe A | **A** | 158 | 692 | 486 | 63 | 115 | 53 | 2 | 2 | 8 | 7 | 10 | 1 | 9 | **1606** |
| Beta-lactam classe B | **B** | 98 | 659 | 2159 | 145 | 81 | 76 | 5 | 5 | 36 | 24 | 46 | 5 | 20 | **3359** |
| Beta-lactam classe C | **C** | 2 | 7 | 15 | 2 | 1 | 0 | 0 | 0 | 0 | 0 | 0 | 0 | 0 | **27** |
| Beta-lactam classe D | **D** | 8 | 499 | 183 | 29 | 10 | 19 | 0 | 1 | 2 | 3 | 9 | 1 | 3 | **767** |
| Aminoglycoside acetyltransferases | **aac** | 112 | 512 | 930 | 51 | 62 | 89 | 3 | 10 | 25 | 25 | 12 | 0 | 0 | **1831** |
| Aminoglycoside adenylyltransferase | **aad** | 1 | 38 | 12 | 0 | 2 | 0 | 0 | 0 | 1 | 0 | 0 | 0 | 0 | **54** |
| Aminoglycoside nucleotidyltransferases | **ant** | 1 | 21 | 19 | 1 | 20 | 1 | 0 | 0 | 1 | 1 | 3 | 0 | 0 | **68** |
| Aminoglycoside phosphotransferases | **aph** | 13 | 104 | 150 | 10 | 8 | 5 | 1 | 1 | 0 | 6 | 4 | 0 | 0 | **302** |
| apramycin resistance gene | **apr** | 0 | 0 | 1 | 0 | 0 | 0 | 0 | 0 | 0 | 0 | 0 | 0 | 0 | **1** |
| Gentamicin acetyltransferase | **gna** | 32 | 328 | 312 | 25 | 16 | 15 | 1 | 2 | 9 | 3 | 3 | 2 | 1 | **749** |
| apramycin resistance gene | **apm** | 0 | 2 | 2 | 0 | 0 | 0 | 0 | 0 | 0 | 0 | 0 | 0 | 0 | **4** |
| streptothricin acetyltransferase | **sat** | 0 | 9 | 4 | 1 | 1 | 0 | 0 | 0 | 0 | 0 | 0 | 0 | 0 | **15** |
| Aminoglycoside resistance methylase | **arm** | 0 | 2 | 0 | 0 | 1 | 0 | 0 | 0 | 0 | 0 | 1 | 0 | 0 | **4** |
| gentamicin resistance methyltransferase | **grm** | 1 | 0 | 3 | 0 | 2 | 0 | 0 | 0 | 0 | 0 | 0 | 0 | 0 | **6** |
| Aminoglycoside resistance methyltransferase | **rmt** | 0 | 1 | 7 | 0 | 4 | 0 | 0 | 0 | 0 | 0 | 0 | 0 | 0 | **12** |
| Sisomicin-gentamicin aminoglycoside resistance methyltransferase | **sgm** | 0 | 0 | 3 | 0 | 6 | 0 | 0 | 0 | 0 | 0 | 0 | 0 | 0 | **9** |
| fosfomycin phosphorylation | **fomA** | 1 | 89 | 21 | 4 | 5 | 0 | 0 | 0 | 0 | 0 | 1 | 0 | 1 | **122** |
| fosfomycin phosphorylation | **fomC** | 0 | 13 | 30 | 1 | 0 | 5 | 0 | 0 | 0 | 0 | 0 | 1 | 0 | **50** |
| fosfomycin resistance methalothiol transferase | **FosA** | 5 | 20 | 21 | 11 | 0 | 2 | 0 | 0 | 0 | 1 | 0 | 0 | 0 | **60** |
| fosfomycin resistance methalothiol transferase | **FosB** | 0 | 34 | 34 | 5 | 6 | 1 | 2 | 0 | 0 | 0 | 4 | 0 | 0 | **86** |
| fosfomycin resistance kinase | **FosC** | 0 | 17 | 10 | 2 | 2 | 0 | 0 | 0 | 0 | 0 | 0 | 0 | 0 | **31** |
| fosfomycin resistance methalothiol transferase | **FosG/FosC** | 2 | 4 | 14 | 2 | 0 | 0 | 0 | 0 | 0 | 0 | 0 | 0 | 0 | **22** |
| fosfomycin resistance hydrolase | **FosX** | 0 | 5 | 3 | 0 | 0 | 0 | 0 | 0 | 0 | 0 | 0 | 0 | 0 | **8** |
| glutathione transferase | **others** | 9 | 94 | 108 | 6 | 3 | 1 | 0 | 0 | 0 | 0 | 0 | 0 | 0 | **221** |
| D-alanine--(R)-lactate ligase VanA | **vanA** | 5 | 52 | 161 | 13 | 5 | 1 | 0 | 0 | 1 | 1 | 0 | 0 | 0 | **239** |
| D-alanine--(R)-lactate ligase VanB | **vanB** | 9 | 62 | 122 | 14 | 4 | 1 | 0 | 1 | 2 | 0 | 1 | 0 | 0 | **216** |
| D-alanine--D-serine ligase VanC1 | **vanC** | 108 | 312 | 516 | 30 | 21 | 32 | 3 | 1 | 18 | 19 | 4 | 0 | 0 | **1064** |
| D-alanine--(R)-lactate ligase VanD | **vanD** | 12 | 128 | 174 | 11 | 17 | 10 | 0 | 1 | 3 | 0 | 4 | 0 | 0 | **360** |
| D-alanine--D-serine ligase VanE | **vanE** | 1 | 10 | 23 | 0 | 0 | 3 | 0 | 0 | 0 | 0 | 0 | 0 | 0 | **37** |
| D-alanine--(R)-lactate ligase VanF | **vanF** | 0 | 18 | 9 | 2 | 0 | 0 | 1 | 0 | 1 | 1 | 0 | 0 | 0 | **32** |
| D-alanine--D-serine ligase VanG | **vanG** | 25 | 77 | 302 | 12 | 2 | 5 | 0 | 0 | 4 | 10 | 5 | 0 | 0 | **442** |
| D-lactate dehydrogenase VanH | **vanH** | 78 | 734 | 1303 | 82 | 108 | 99 | 2 | 3 | 32 | 17 | 14 | 3 | 0 | **2475** |
| D-alanine--(R)-lactate ligase VanI | **vanI** | 2 | 25 | 54 | 1 | 4 | 2 | 0 | 0 | 0 | 0 | 1 | 0 | 0 | **89** |
| peptidoglycan bridge formation peptidyltransferase VanK-I | **vanK** | 185 | 706 | 796 | 61 | 90 | 80 | 1 | 2 | 31 | 30 | 18 | 2 | 9 | **2011** |
| D-alanine--D-serine ligase VanL | **vanL** | 15 | 34 | 69 | 4 | 5 | 2 | 0 | 0 | 0 | 0 | 1 | 0 | 0 | **130** |
| D-alanine--(R)-lactate ligase VanM | **vanM** | 1 | 11 | 25 | 2 | 0 | 0 | 0 | 0 | 0 | 0 | 0 | 0 | 0 | **39** |
| D-alanine--D-serine ligase VanN | **vanN** | 2 | 21 | 30 | 0 | 3 | 3 | 0 | 0 | 2 | 0 | 0 | 0 | 0 | **61** |
| D-alanine--(R)-lactate ligase VanO | **vanO** | 3 | 9 | 10 | 1 | 0 | 0 | 1 | 0 | 0 | 0 | 1 | 0 | 0 | **25** |
| VanA-type vancomycin resistance histidine kinase VanS | **vanS** | 125 | 170 | 517 | 25 | 40 | 37 | 4 | 2 | 8 | 17 | 4 | 1 | 0 | **950** |
| membrane-bound serine racemase VanT-C | **vanT** | 25 | 147 | 652 | 41 | 7 | 30 | 0 | 3 | 21 | 25 | 7 | 0 | 10 | **968** |
| glycopeptide resistance accessory protein VanW-B | **vanW** | 0 | 215 | 202 | 13 | 12 | 25 | 0 | 1 | 5 | 27 | 0 | 0 | 2 | **502** |
| D-Ala-D-Ala dipeptidase VanX-A | **vanX** | 66 | 51 | 128 | 8 | 1 | 11 | 0 | 1 | 0 | 7 | 9 | 0 | 0 | **282** |
| D-Ala-D-Ala carboxypeptidase VanY-A | **vanY** | 85 | 28 | 308 | 24 | 16 | 2 | 2 | 1 | 0 | 0 | 29 | 0 | 0 | **495** |
| glycopeptide resistance protein VanZ-A | **vanZ** | 7 | 2 | 11 | 1 | 0 | 0 | 1 | 0 | 0 | 0 | 0 | 0 | 0 | **22** |
| 23S rRNA (guanine(748)-N(1))-methyltransferase ChrB | **chr** | 1 | 19 | 23 | 5 | 0 | 0 | 0 | 1 | 2 | 0 | 0 | 0 | 0 | **51** |
| rRNA methylase | **emt** | 2 | 22 | 47 | 8 | 1 | 5 | 0 | 0 | 2 | 0 | 14 | 0 | 0 | **101** |
| inactive erythromycin esterase | **ere** | 1 | 0 | 5 | 1 | 0 | 0 | 0 | 0 | 0 | 0 | 0 | 0 | 0 | **7** |
| lipophilic membrane protein | **llm** | 2 | 65 | 174 | 11 | 2 | 14 | 0 | 0 | 0 | 3 | 2 | 0 | 0 | **273** |
| Lincosamide nucleotidyltransferase | **lnu** | 8 | 40 | 10 | 1 | 0 | 1 | 0 | 1 | 1 | 1 | 0 | 0 | 0 | **63** |
| macrolide-inactivating glycosyltransferase | **mgt** | 1 | 6 | 1 | 0 | 0 | 0 | 0 | 0 | 0 | 0 | 0 | 0 | 0 | **8** |
| macrolide phosphotransferase | **mph** | 2 | 7 | 18 | 0 | 0 | 0 | 0 | 0 | 0 | 0 | 1 | 0 | 0 | **28** |
| Mycinamicin resistance protein homolog | **myr** | 0 | 2 | 4 | 1 | 0 | 0 | 0 | 0 | 0 | 0 | 0 | 0 | 0 | **7** |
| ABC-F type ribosomal protection protein | **ole** | 2 | 12 | 1 | 0 | 0 | 1 | 0 | 0 | 0 | 0 | 1 | 0 | 0 | **17** |
| streptogramin A O-acetyltransferase | **vat** | 9 | 220 | 68 | 8 | 5 | 9 | 6 | 2 | 4 | 0 | 6 | 0 | 1 | **338** |
| streptogramin B lyase | **vgb** | 11 | 3 | 10 | 0 | 0 | 0 | 0 | 0 | 0 | 0 | 0 | 0 | 0 | **24** |
|  | **vgh** | 1 | 1 | 4 | 0 | 0 | 0 | 0 | 0 | 0 | 0 | 0 | 0 | 0 | **6** |
| 23S rRNA (adenine(2503)-C(8))-methyltransferase Cfr | **cfr** | 2 | 64 | 438 | 38 | 14 | 43 | 16 | 7 | 0 | 0 | 26 | 0 | 0 | **648** |
| erythromycin esterase | **erm** | 338 | 273 | 1902 | 125 | 166 | 115 | 7 | 5 | 47 | 34 | 51 | 2 | 12 | **3077** |
|  | **mls** | 0 | 0 | 1 | 0 | 0 | 0 | 0 | 0 | 0 | 0 | 0 | 0 | 0 | **1** |
| Methyltransferase R mycinamicin, tylosin and lincosamides | **rlm** | 6 | 133 | 100 | 7 | 3 | 10 | 0 | 0 | 1 | 2 | 0 | 0 | 1 | **263** |
| 23S rRNA (adenine(2503)-C(8))-methyltransferase CipA | **cip** | 2 | 4 | 90 | 3 | 7 | 23 | 3 | 0 | 0 | 0 | 2 | 1 | 0 | **135** |
| clbC is a plasmid-encoded cfr gene | **clb** | 0 | 4 | 61 | 2 | 0 | 12 | 1 | 1 | 1 | 0 | 2 | 0 | 0 | **84** |
| Chloramphenicol acetyltransferase | **cat** | 5 | 402 | 86 | 8 | 10 | 24 | 0 | 0 | 6 | 0 | 5 | 0 | 0 | **546** |
| chloramphenicol phosphotransferase | **cpt** | 1 | 0 | 0 | 0 | 1 | 0 | 0 | 0 | 0 | 0 | 1 | 0 | 0 | **3** |
| quinolone resistance genes | **qnr** | 23 | 69 | 104 | 16 | 8 | 10 | 0 | 1 | 1 | 3 | 4 | 0 | 0 | **239** |
| rifampin monooxygenase Rox | **rox** | 0 | 0 | 7 | 1 | 0 | 1 | 0 | 0 | 0 | 0 | 0 | 0 | 0 | **9** |
| rifampin ADP-ribosyltransferase | **arr** | 1 | 0 | 8 | 0 | 0 | 6 | 0 | 0 | 0 | 0 | 0 | 0 | 0 | **15** |
| rifampin monooxygenase Iri | **iri** | 0 | 0 | 7 | 1 | 0 | 0 | 0 | 0 | 0 | 0 | 0 | 0 | 0 | **8** |
| Rifampin phosphotransferase | **rph** | 1 | 100 | 40 | 2 | 9 | 0 | 0 | 0 | 4 | 0 | 1 | 0 | 0 | **157** |
| tetracycline-inactivating monooxygenase Tet(X1) | **tetX** | 1 | 4 | 5 | 1 | 0 | 7 | 0 | 0 | 0 | 0 | 0 | 0 | 0 | **18** |
| oxytetracycline resistance phosphoribosyltransferase domain-containing protein Tet(34) | **tet34** | 5 | 54 | 41 | 3 | 1 | 0 | 0 | 1 | 3 | 0 | 0 | 0 | 1 | **109** |
| tetracycline resistance ribosomal protection protein Otr(A) | **otrA** | 0 | 21 | 49 | 3 | 0 | 0 | 2 | 1 | 0 | 1 | 1 | 0 | 0 | **78** |
| tetracycline resistance ribosomal protection protein Tet(32) | **tet32** | 8 | 7 | 37 | 1 | 3 | 1 | 9 | 0 | 0 | 0 | 0 | 0 | 0 | **66** |
| tetracycline resistance ribosomal protection protein Tet(36) | **tet36** | 3 | 3 | 31 | 4 | 0 | 0 | 0 | 3 | 2 | 2 | 0 | 0 | 0 | **48** |
| tetracycline resistance ribosomal protection protein Tet(44) | **tet44** | 22 | 15 | 50 | 3 | 0 | 3 | 2 | 0 | 2 | 0 | 2 | 0 | 0 | **99** |
| tetracycline resistance ribosomal protection protein Tet(BP) | **tetBP** | 26 | 147 | 475 | 35 | 25 | 17 | 11 | 2 | 10 | 1 | 25 | 0 | 4 | **778** |
| tetracycline resistance ribosomal protection protein Tet(M) | **tetM** | 16 | 55 | 185 | 8 | 1 | 1 | 1 | 0 | 4 | 0 | 3 | 0 | 0 | **274** |
| tetracycline resistance ribosomal protection protein Tet(O) | **tetO** | 4 | 20 | 22 | 4 | 0 | 0 | 0 | 0 | 0 | 1 | 1 | 0 | 0 | **52** |
| tetracycline resistance ribosomal protection protein | **TetO/32** | 7 | 1 | 6 | 0 | 0 | 1 | 0 | 0 | 2 | 0 | 0 | 0 | 0 | **17** |
| tetracycline resistance ribosomal protection protein | **tetQ** | 12 | 14 | 23 | 5 | 0 | 3 | 1 | 1 | 0 | 0 | 1 | 0 | 0 | **60** |
| tetracycline resistance ribosomal protection protein | **tetS** | 2 | 8 | 9 | 2 | 0 | 1 | 0 | 2 | 1 | 0 | 0 | 1 | 0 | **26** |
| tetracycline resistance ribosomal protection protein | **tetT** | 177 | 467 | 1244 | 85 | 96 | 79 | 11 | 5 | 26 | 30 | 16 | 1 | 6 | **2243** |
| tetracycline resistance ribosomal protection protein | **tetW** | 29 | 200 | 252 | 20 | 21 | 25 | 1 | 0 | 8 | 4 | 3 | 0 | 1 | **564** |
| pyrazinamidase | **Pnc** | 18 | 20 | 52 | 2 | 1 | 0 | 0 | 1 | 0 | 0 | 1 | 0 | 0 | **95** |
| Nitroimidazole resistance genes | **nim** | 10 | 7 | 5 | 0 | 0 | 16 | 1 | 0 | 0 | 0 | 0 | 0 | 0 | **39** |
| Bah amidohydrolases are membrane proteins that inactivate bacitracin. | **bah** | 0 | 0 | 1 | 0 | 0 | 0 | 0 | 0 | 0 | 0 | 0 | 0 | 0 | **1** |
| intrinsic colistin resistance enzyme | **icr** | 0 | 0 | 1 | 0 | 0 | 0 | 0 | 0 | 0 | 0 | 0 | 0 | 0 | **1** |
| mobilized colistin resistance gene | **mcr** | 2 | 4 | 8 | 1 | 0 | 0 | 0 | 0 | 0 | 0 | 0 | 0 | 0 | **15** |
| fusidic acid resistance | **fus** | 0 | 0 | 1 | 0 | 0 | 0 | 0 | 0 | 0 | 0 | 0 | 0 | 0 | **1** |
|  | **Total** | **1958** | **8455** | **15645** | **1118** | **1054** | **1038** | **101** | **71** | **337** | **306** | **360** | **21** | **81** | **30545** |
